# 25-Hydroxy Cholesterol Effectively Inhibits Japanese Encephalitis Virus Infection in a Cellular Model

**DOI:** 10.1101/2024.12.18.628890

**Authors:** Km. Archana, Bushra Qazi, Babita Bohra, Raj Kamal Tripathi, Sourav Haldar

**Affiliations:** Division of Virus Research and Therapeutics, CSIR-Central Drug Research Institute, Lucknow (226031), UP, India; Academy of Scientific and Innovative Research (AcSIR), Ghaziabad 201002, India

**Author notes:** Corresponding Authors: Sourav Haldar,) and Rajkamal Tripathi.

**Keywords:** Japanese encephalitis virus, 25-hydroxycholesterol, flavivirus, membranes, cholesterol

## Abstract

The Japanese encephalitis virus (JEV) is a mosquito-borne, enveloped flavivirus that causes acute encephalitis. Although JEV is increasingly recognized as a global threat, there is currently no FDA-approved treatment available for JEV. 25-hydroxycholesterol (25-HC) is an oxysterol produced through the oxidation of cholesterol, a process catalyzed by cholesterol 25-hydroxylase (CH25H), which is an interferon-stimulated gene that is upregulated during viral infections. In this study, we report for the first time that 25-HC effectively prevents JEV infection in cells. Our results show that 25-HC inhibits JEV infection in Vero cells with an IC50 of ∼ 4.18 µM, while displaying no cell toxicity (CC50 > 50 µM). In addition, we show that RNA level of the JEV envelope (E) protein exhibits a dose-dependent reduction in the presence of 25 HC, which is further supported by the concomitant decrease of the E protein expression. We surmise that the anti-JEV properties of 25 HC stem from its effects on the host plasma membrane organization and dynamics by modulating membrane cholesterol level, which may impair the entry and fusion of JEV. Given the antiviral effects of 25HC against Zika and Dengue viruses, our findings suggest that 25-HC could be a promising candidate for developing broad-spectrum anti-flaviviral therapeutics.

## Introduction

The Japanese encephalitis virus (JEV) is a member of the Flaviviridae family that is transmitted via mosquitoes. JEV mainly infects various cell types in the CNS and causes acute encephalitis. Its primary distribution areas are tropical parts of Asia, such as Japan, China, Taiwan, Korea, the Philippines, India, and all of Southeastern Asia. JEV is becoming more widespread throughout Asia due to the increase in mosquito population and is also rapidly spreading worldwide. More than 50,000 cases of JEV infection with a 20–30% mortality are reported annually. Most of the JEV-infected cases remain asymptomatic. Survivors suffer neurological or neuropsychiatric after effects that mimic Parkinsonian movement disorders, paralysis similar to that of poliomyelitis, or cognitive impairment.^1^ Children under the age of 14 are most commonly affected by the disease. However, there are no FDA-approved drugs against JEV and available vaccines are not completely effective.^2^

JEV is an enveloped, positive-sense, single-stranded RNA virus with a diameter of around 50 nm with spheroid cubical symmetry. Its 11000 nucleotides genome is translated in a single polyprotein precursor, which is then cleaved into three structural [Capsid(C), pre-membrane (prM), and Envelope(E)] and seven nonstructural (NS1, NS2A, NS2B, NS3, NS4A, NS4B, and NS5) proteins by host and viral proteases.^3^ Receptor-mediated endocytosis and low pH-triggered membrane fusion within endosomes allow JEV to transfer their genome inside the host cells.^4^ Subsequently newly synthesized viral components are assembled and packaged at the host endoplasmic reticulum (ER) membrane and released by exocytosis. Host membranes play crucial roles in different stages of the JEV infection, particularly during the entry processes.^5^

25-hydroxycholesterol (25-HC) is an oxysterol that is produced by oxidation of cholesterol catalyzed by cholesterol 25-hydroxylase (CH25H). CH25H is an interferon-stimulated gene that is upregulated upon viral infection. ^6^ CH25H and its product 25HC regulate the cholesterol content of membranes in different subcellular compartments. 25HC promotes cholesterol trafficking into the endoplasmic reticulum^7^, and 25 HC treatment decreases plasma membrane cholesterol level^8^ by increasing the accessibility of membrane cholesterol and inducing lipid droplet formation.^9^ Moreover, 25 HC alters the plasma membrane properties such as domain organizations and fluidity. ^10^ 25-HC treatment leads to antiviral activity against a wide range of viruses such as SARS CoV2, Zika, Ebola virus (EBOV), Nipah virus (NiV), and human immunodeficiency virus (HIV). On the other hand, 25 HC does not affect Adeno virus infection.^6^ Although the antiviral effect of 25 HC against Zika and Dengue have been reported ^6^, the antiviral effect of 25HC against JEV has not been explored. Since plasma membrane cholesterol plays an important role in the entry of JEV, ^5^ and 25 HC modulates cholesterol organization in plasma membranes, we hypothesize that 25HC treatment will inhibit JEV infection. Our results show that 25 HC is a potent inhibitor of JEV infection in cells.

## Materials and Methods

### Cytotoxicity Assay

Vero Cells (Vero Cells C1008, cat-CRL-1586 ATCC) were cultured in growth medium [DMEM media (D5648-L, Sigma) with 10% FBS (10270-106, GIBCO)] under 5% CO_2_ at 37°C. Cells were seeded in a 96well cell culture plate (1*10^4^/well) in 200 μl/well. After 12 hours of seeding, 25-Hydroxycholesterol (H1015-25MG, SIGMA) was added at different concentrations in triplicates with vehicle (DMSO) control. The plate was incubated at 37°C with 5% CO_2_. At 68 hrs, MTT (3-(4,5-Dimethylthiazol-2-yl)-2,5-Diphenyltetrazolium Bromide, CAS298-93-1, Sigma) was added. The plate was incubated at 37 °C with 5% CO_2_. At 72 hrs, the supernatant was removed carefully and DMSO (100 μl/well) was added to dissolve crystals. The plate was kept at shaking condition for 10-15 minutes. Absorbance was measured at 570 nm using a microplate reader.

### Flow Cytometry

Vero cells were seeded in a 96 well plate. Cells were pretreated with 25-HC at different concentrations before 14hrs of infection followed by infection with JE virus [Japanese Encephalitis Virus (Vellore Strain) was a kind gift from Dr Vikas Agrawal at the Sanjay Gandhi Postgraduate Institute of Medical Sciences (SGPGIMS), Lucknow] at .01 MOI. The plate was incubated for 2 hrs under humid conditions in 5%CO_2_. At 2 hrs post infection, infection media was removed and fresh media at 200µl/well (DMEM with 2 %FBS) with 25 HC (at the same concentrations of the pre-treatment) was added. The plate was incubated under humid condition in 5% CO_2_ for 72 hrs. Overlay media was aspirated and cells were washed with 1x PBS, 150 µl/well. Cells were harvested by adding 0.25% Trypsin-EDTA (25 µl/well). Next, cells were transferred to a U-bottom 96 well plate and were fixed with 4% paraformaldehyde (50 µl/well). After proper mixing, and the plate was incubated for 10 mins at RT to fix the cells. Cells were permeabilized with 1x permeabilization buffer at 150 µl/well (2%BSA+ 0.1%Saponin in PBS). Next, 40 µl/well blocking solution (1% normal mouse serum in 1x permeabilization buffer) was added. After 30 minutes incubation at RT, 4G2-alexa488 mAb (20 µl/well) was added, antibody dilution was prepared in blocking buffer (1:400). The Plate was incubated at 37°C with shaking. After that the cells were washed with 1X permeabilization and buffer, and pelleted by centrifugation at 1000g. The cell pellet was resuspended in 1x PBS (200 µl/well). The samples were read in a flow cytometer (BD Bioscience-FACS LYRIC). Data was analyzed using FlowJo (BD Bioscience).

### Confocal Microscopy

Cells were seeded on coverslips and treated with 25-HC for 24 hours. The cell monolayer was washed with 1X PBS and was fixed with 4% PFA. Next, cells were permeabilized using TritonX100 (0.1%) followed by washing with PBS. Cells were stained with Nile red (N30/3-1G, sigma) by incubating with Nile red. After that the cells were washed with PBS. The coverslips were mounted on a glass slide with a mounting media containing DAPI (F6057, Sigma). The slides were imaged using an Olympus confocal microscope.

### Antigen Release Assay

Vero Cells were cultured in a growth under 5% CO_2_ at 37°C. After 14 hours of seeding cells were pretreated with 25-HC at different concentrations (20μM, 10μM, 5μM, 2.5μM, and 1.25μM). After 14 hrs, Cells were infected with the Japanese Encephalitis Virus at 0.01 MOI. 2% FBS was used in infection media. The plate was incubated at 37°C with 5% CO_2_, during incubation plate was Shaken at 30-minute intervals to avoid drying. At 2 hours postinfection, the infection media with virus was replaced with fresh DMEM containing 2% FBS along with varying concentrations of 25HC. At 72 hrs post-infection, the supernatant was collected to check viral (antigen) release in the supernatant using ELISA. 96 well, high binding ELISA Plate (Corning-3590) was coated with 100μl/well coating buffer [0.1M Sodium Bicarbonate solution, pH-9.6 with 2.5 µg/ml 4G2 antibody (anti-flavivirus group antigen antibody 4G2).^11^ The plate was incubated overnight at 4°C, followed by three times 1XPBS washing. For blocking, 5% skim milk in 1xPBS was used. The plate was again washed with 1x PBS three times (150 μl/well). Each sample was added at 100 μl/well in triplicates. The plate was incubated at 37°C for one hour followed by washing with PBST (0.1% tween20). HRP-tagged 4G2 antibody (1:4000),100 μl/well was added. After one hour of incubation, three times PBST washing was done. TMB substrate (TMBW-0100-01, Surmodics) was added (100 μl/well) in the dark. The plate was incubated at 37°C for 30 minutes. In the end, 7% (v/v) H_2_SO_4_ was added (50 μl/well). The plate was read at 450 nm in an ELISA plate Reader in chemiluminescence mode.

### Western Blot Analysis

Cells were seeded in growth medium and pretreated with 25-HC at test concentrations (1.25μM-20μM). After 14 hrs, Cells were infected with the Japanese Encephalitis Virus at 0.01 MOI. 2% FBS was used in the infection media. The plate was incubated at 37°C with 5% CO_2_, During incubation, the plate was shaken at 30-minute intervals to avoid drying. At 2 hours postinfection, the infection media with virus was replaced with fresh DMEM containing 2% FBS along with varying concentrations of 25HC, at 72 hrs post-infection. After removing the media, cells were washed with ice-cold PBS and scraped in a lysis buffer (RIPA-HIMEDIA-TCL131) in the presence of a protease inhibitor cocktail (SIGMA-100XP8340). To prepare cell lysate, cells were subjected to sonication for 90 seconds (pulse 10, Amp1 40%) (Fisher Scientific) and then pelleted by centrifugation at 16000 RCF for 20 minutes at 4°C. The supernatant was collected and stored at −20°C. Total protein concentration was determined by a bicinchoninic acid assay, using a kit (Thermofisher, USA BCA-23227). Protein lysates were separated on an SDS-PAGE (10%) and then transferred to a PVDF membrane (Bio-Rad-1620177). The membrane was blocked in 5% skimmed milk (Bio-Rad-1706404) in PBS containing 0.1% Tween 20 (Amresco-0777-1L) for 1 hour with slow rocking at room temperature. JEV Envelope (E) protein was detected by a polyclonal rabbit anti-E antibody [Invitrogen-PA5-111964(1:10,000)] as primary, and an HRP conjugated goat anti-rabbit IgG H&L[ab6721 (1:10,000)] was used as a secondary antibody. The same blots were used after stripping for estimating β-actin as a loading control using an HRP-conjugated primary antibody [Abcam 8226 (1:10,000)]. Quantification of Western blots was done using ImageJ software (NIH, Bethesda, Maryland, USA). The band intensity values obtained were divided by the corresponding β-actin loading control.

### RT-qPCR analysis

Vero cells were seeded (.25×10^6^cells /well) in a 6-well culture plate. Cells were pre-treated with 25HC at various concentrations (1.25μΜ to 20μM). After 14 hrs of pre-treatment, cells were infected with JEV at .01 MOI. After 2 hrs post-infection, the same concentrations of 25 HC were put back. After 72 hrs post-infection, cells were collected. Total RNA isolation was done using the TRIzol reagent (Thermo Fisher Scientific). cDNA synthesis was done with the Verso cDNA Synthesis Kit (Thermo Fisher Scientific AB-1453/A) using C1000 Touch Thermal Cycler (Bio-Rad). 2μg of RNA was used for cDNA synthesis. RT-qPCR was done in CFX96 Real Time PCR system (Bio-Rad) using Bio-Rad universal Syber Green mastermix (1725120) using 500ng of cDNA. For RT-qPCR, primers were designed and purchased from IDT for two structural genes (Envelope and Capsid) of JEV; [C(FP)-5’ GTGGGAGTGAAGAGGGTAGTAA 3’, C(RP)-5’ CTGCTTTCCATCGGCCTAAA 3’ (133bp), E(FP)-5’ TGCATGGAACCACCACTT3’, E(RP)-5’ GAAGGAGCATTGGGTGTTATTG 3’ (97bp)]. Statistical analysis of the results was carried out using one-way ANOVA analysis and normalized to GAPDH.

### Results and Discussion

We wanted to investigate whether 25 HC inhibits JEV infection in cells. We used Vero cells for this purpose which is used to study JEV infection^12^. First, we evaluated the cytotoxicity of 25HC in Vero cells by the MTT assay. We did not observe any significant change in cell viability in the presence of 25 HC up to 50 µM in 72 hours (Figure 1 (A)). Next, we evaluated the effect of 25 HC on the infectivity of JEV in Vero cells by estimating the intracellular viral load by FACS and extracellular release of viral antigen in the supernatant by ELISA. We incubated Vero cells with varying amounts of 25 HC for 14 hours before infection according to previous reports.^6^ Figure 2 shows Alexa 488-4G2 positive cells in the scatter plot as a function of 25 HC concentration. Flow cytometry data reveals that the intracellular viral load decreased by about 87% in 25-HC pre and post-treated Vero cells at 20μM with an IC_50_ of 4.18 ± 0.75 μM (SEM, n = 3) (Figure 1 (B)). Next, we wanted to evaluate whether 25HC has any effect on virus release. For this, we determined the level of viral antigen (the Envelope protein) released in the media in the presence of varying amounts of 25 HC by ELISA (See Figure 2 A).

**Figure 1.**
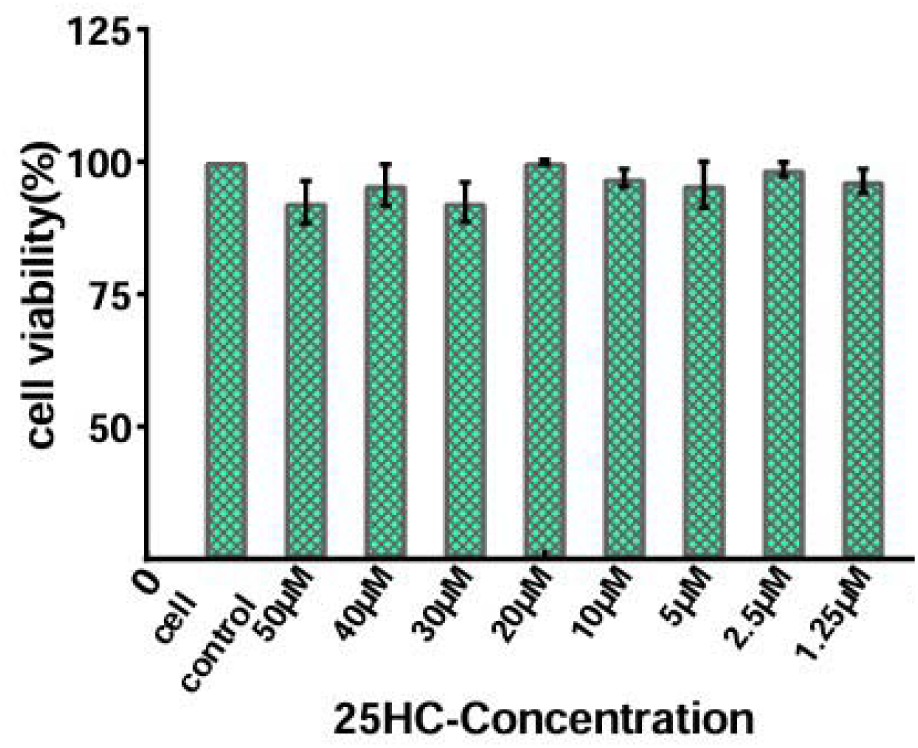
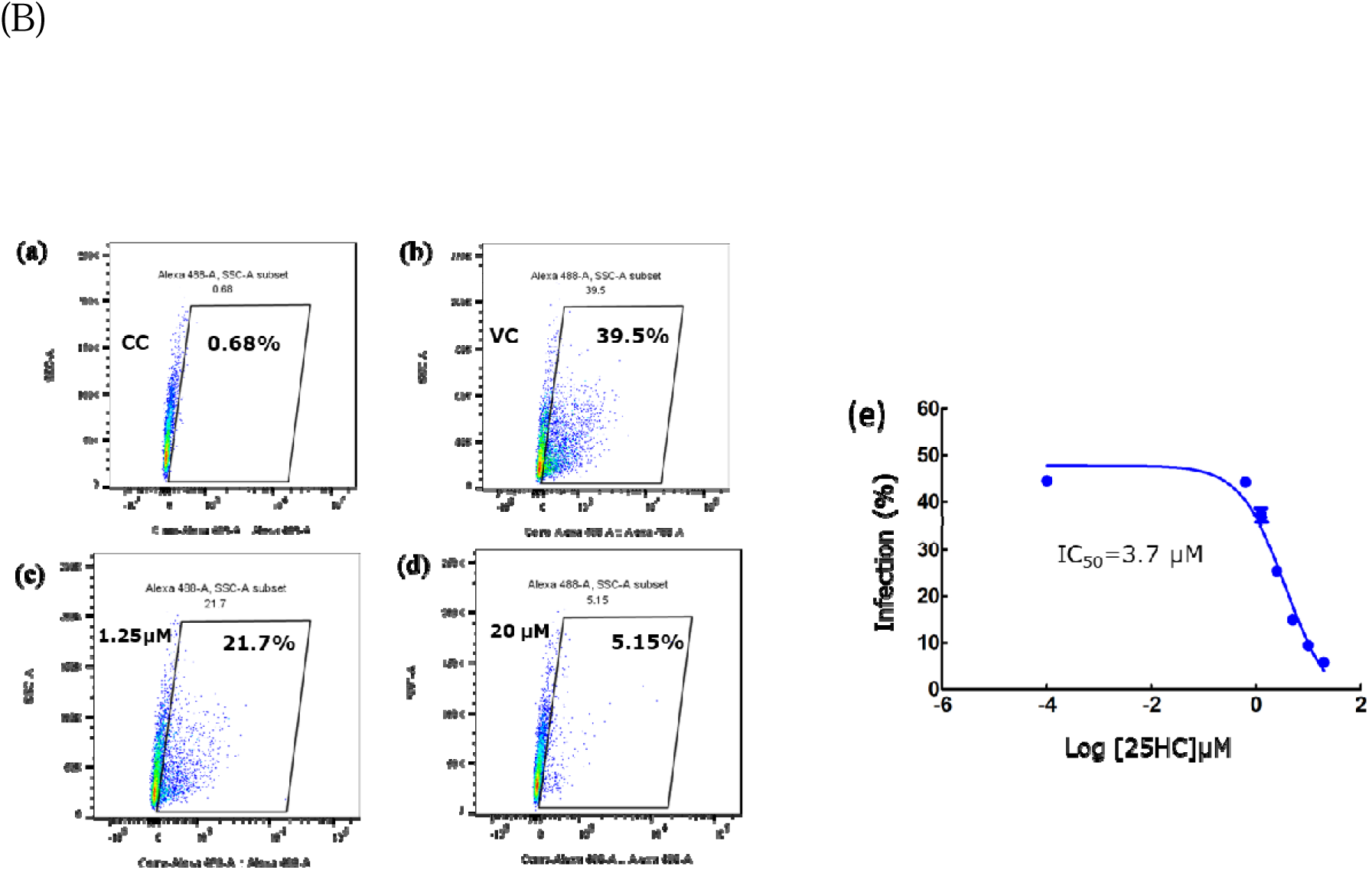
(A) Cell viability as a function of 25 HC concentration. Vero cells were incubated with increasing amounts of 25 HC in DMSO (up to 50 µM) for 72 hours. Cell viability was determined by MTT assay. Data represent means ± SD of three replicates. See the methods section for details. (B) Effect of Pre and post-treatment 25HC on JEV infection. Representative flow cell scatter diagrams show the reduction in JEV infections in Vero cells in the presence of 25 HC. Vero cell were pre-treated with varying amounts of 25 HC followed by infection with JEV at 0.01 MOI. Cells were grown for 72 hours in the presence of (same) varying amounts of 25 HC. Cells were stained with Alexa488 labeled 4G2 Antibody against the fusion domain of envelope protein. Numbers indicate the percentage of Alexa 488 + virus-infected cells within the entire population (a) Cell control, (b) Virus control without 25 HC treatment, (c) sample treated with 1.25μM 25HC, (d) sample treated with 20μM 25HC. Data was analyzed using Flowjo software. (e) A dose-response curve showing the effect of 25 HC on JEV infection. Data represent mean ± SD from at least three independent experiments. See the methods section for details.

**Figure 2.**
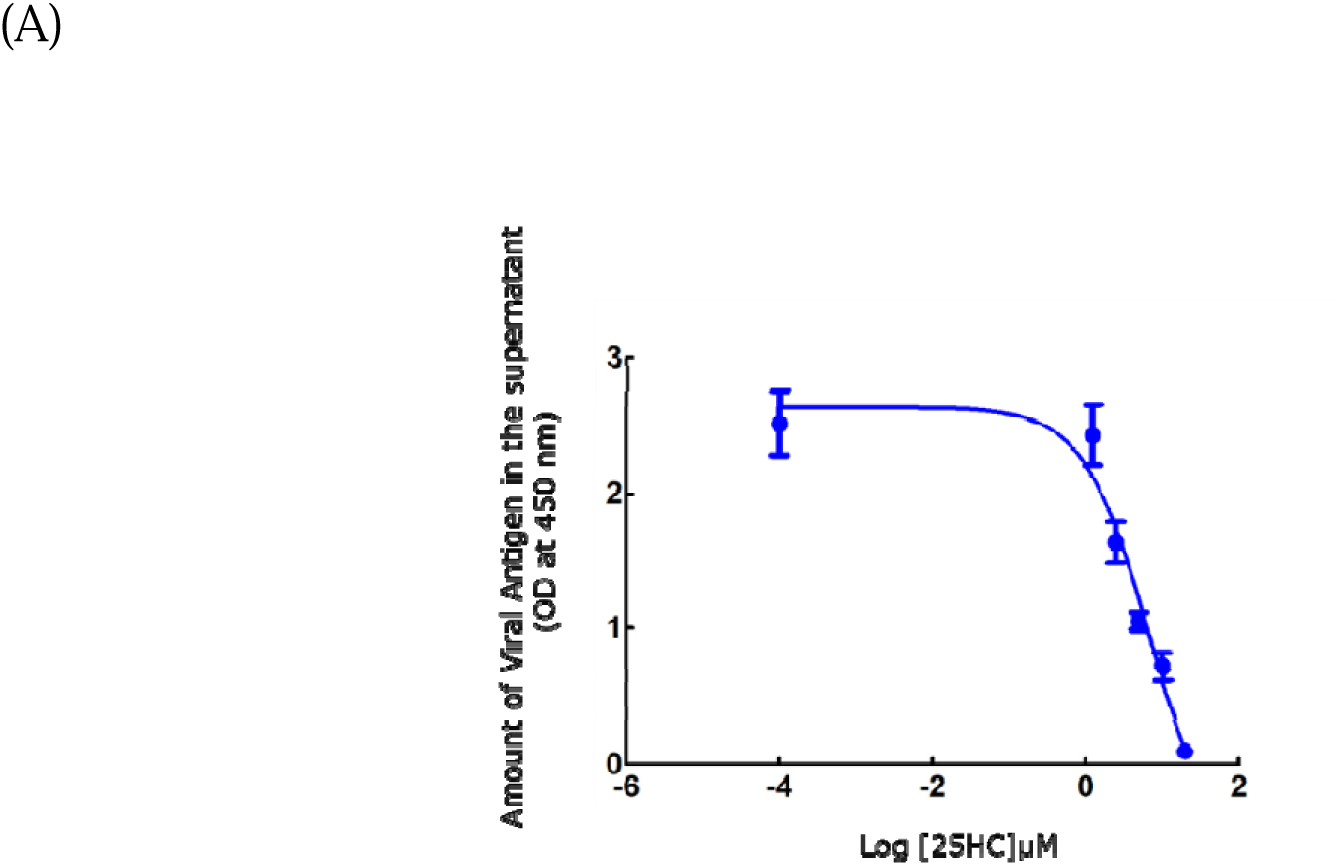
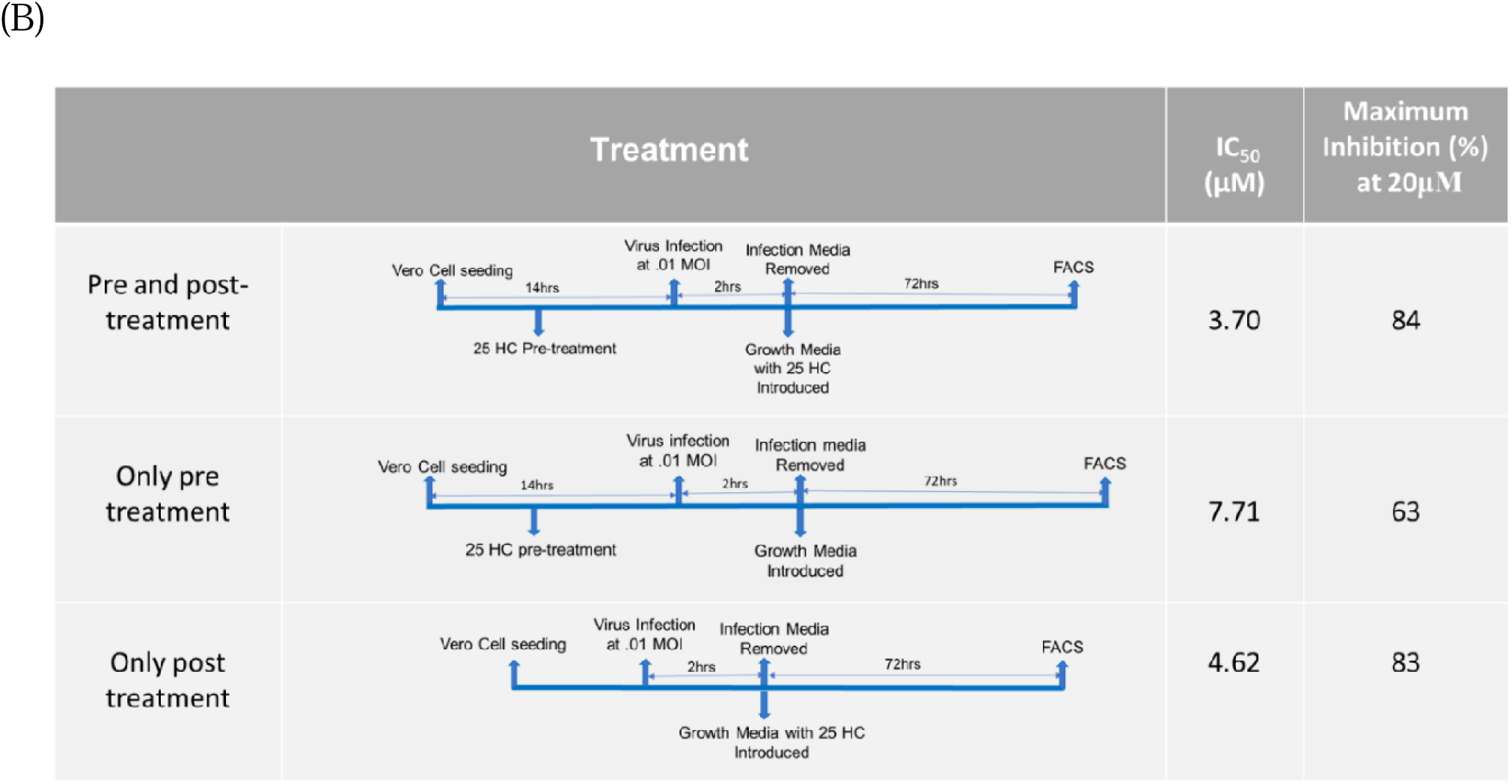
**(A)**. The inhibitory effect of 25HC on JEV release. Vero cells were pre-treated with varying amounts of 25 HC followed by infection with JEV at 0.01 MOI. Cells were grown for 72 hours in the presence of (same) varying amounts of 25 HC. The amount of spike protein released in the media was determined by ELISA using HRP conjugated 4G2 antibody against the envelope protein. The figure shows the amount of antigen as a function of 25 HC. Data represent mean ± SD from at least three independent experiments. See the methods section for details. **(B).** Comparison of the anti-JEV effects of 25 HC under different conditions.

Our results show that 25 HC treatment reduced viral release from Vero cells quite effectively with an IC_50_ of 5.91 ± 0.80 µM (SEM, n=3). There is no significant difference between the IC50 values for infectivity measured by FACS and virus release determined by ELISA. Our data therefore suggest that 25 HC likely influences the early stages of infection, with its effect on viral release being a consequence of its impact on infectivity. In other words, 25 HC may not directly affect the egress of the virus from infected cells. (A)

Next, we wanted to check whether 25HC has any impact on the first round of infection. For this, we monitored intracellular viral load after 12 hours of infection. Our results show that even at early time point 25 HC reduced (relative) infection to ∼ 24 % in the presence of 20 µM 25 HC (See Figure 3).

**Figure 3.**
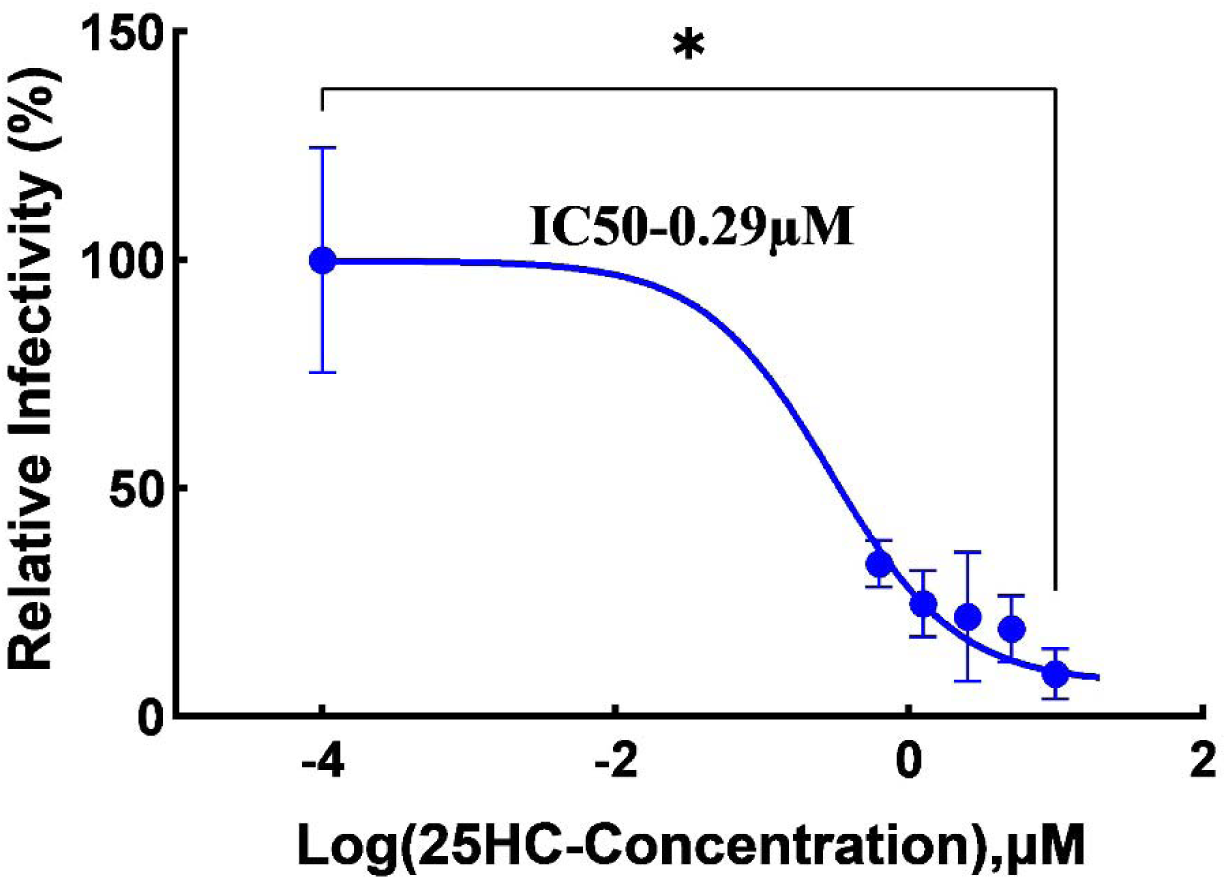
Antiviral effect of JEV at early time point. The figure shows the reduction in infection (determined by FACS) in the presence of 25 HC, assessed after 12 hours of infection. Vero cells were pre-treated with varying amounts of 25 HC followed by infection with JEV at 0.01 MOI. Cells were grown for 12 hours in the presence of (same) varying amounts of 25 HC. Cells were stained with Alexa488 labeled 4G2 Antibody against the envelope protein. Data represent mean ± SD from at least three independent experiments. See the methods section for details (* implies p < 0.05).

We confirmed the inhibition of JEV infection in the presence of 25 HC by determining the reduction of the viral RNA level by RT-qPCR and reduction viral protein expression by western blot analysis. Figure 4(A) shows the dose-dependent decrease of Envelope (E) and Capsid (C) protein RNA levels in the presence of 25 HC by RT qPCR. Figure 4 B shows the concomitant dose-dependent reduction in JEV E-protein expression level in the presence of 25 HC. Taken together, our results show that 25 HC treatment effectively inhibits JEV infection.

**Figure 4.**
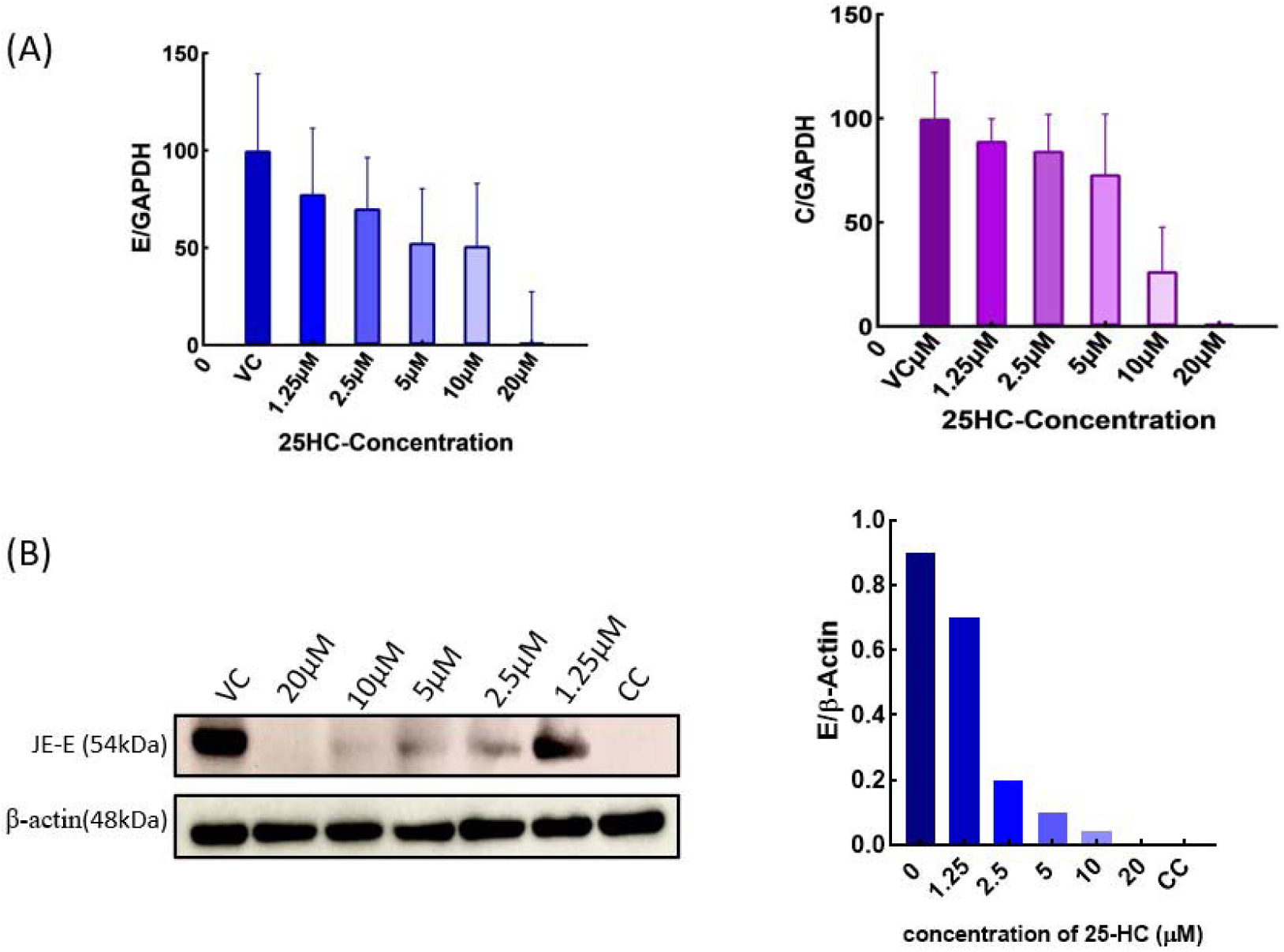
**(A)** Estimation of viral RNA content in the presence of 25HC using RT-qPCR. Concentration-dependent decrease in viral RNA content was observed when cells were incubated with the virus in the presence of 25HC. The viral Envelope (E) and Capsid (C) genes were detected to find the viral load in the presence of 25HC. All the data are represented as mean ± SEM for n = 2. **(B)** Decrease in JEV Envelope € protein expression by Western blot analysis. Cell lysates were prepared from pre- and post-treated with varying amount of 25 HC after 72 hours of infection. β actin was taken as loading control. See the Materials and method sections for details.

To delineate the mechanism of the antiviral effect of 25HC against JEV, we carried out the time of addition assay.^13^ We compared the effect of pre-and post-treatment with 25 HC on the JEV infectivity with the corresponding effect of only pre-treatment and only post-treatment (Figure 2B). Interestingly, only the pre-treatment condition exhibited the highest IC_50_ of 6.2 µM, where pre-and post-treatment yielded an IC_50_ of 3.7 µM, while only post-treatment inhibited infection with an IC_50_ of 4.6 µM (Figure 2B). This suggests that a combination of pre and post-treatment brought about the most effective inhibition and 25 HC might affect multiple stages of the infection, e.g., viral entry and fusion and packaging.

25HC could inhibit viral infection by directly inhibiting endocytosis or by preventing fusion of viral envelope with endosomal membranes. ^14^ It was earlier shown that lipid rafts play a crucial role in JEV entry and replication, and cholesterol depletion by mβCD (disruption of lipid rafts) reduced JEV infection in neuronal cells.^5^ Interestingly, 25 HC is known to alter cholesterol organization in the plasma membranes by increasing cholesterol accessibility and reducing plasma membrane cholesterol. Therefore, it is plausible that 25 HC exerts its antiviral effect against JEV by inhibiting raft-dependent endocytosis via altered cholesterol organization in the plasma membrane. Previously, it was shown that 25HC enhances cholesterol esterification and promotes cholesterol transport from the plasma membrane to the ER.^7^ 25HC depletes accessible cholesterol on the plasma membrane through activating acyl-CoA: cholesterol acyltransferase (ACAT) ^14^ and leads to enhanced lipid droplet formation. To test whether 25HC treatment has any effect on membrane cholesterol level and subsequent lipid reorganization, we checked the level of lipid droplets by Nile red staining upon 25 HC treatment. Our results show (Figure 5) a significant increase in lipid droplet (staining) in the presence of 25 HC, as reported earlier.^15^

**Figure 5:**
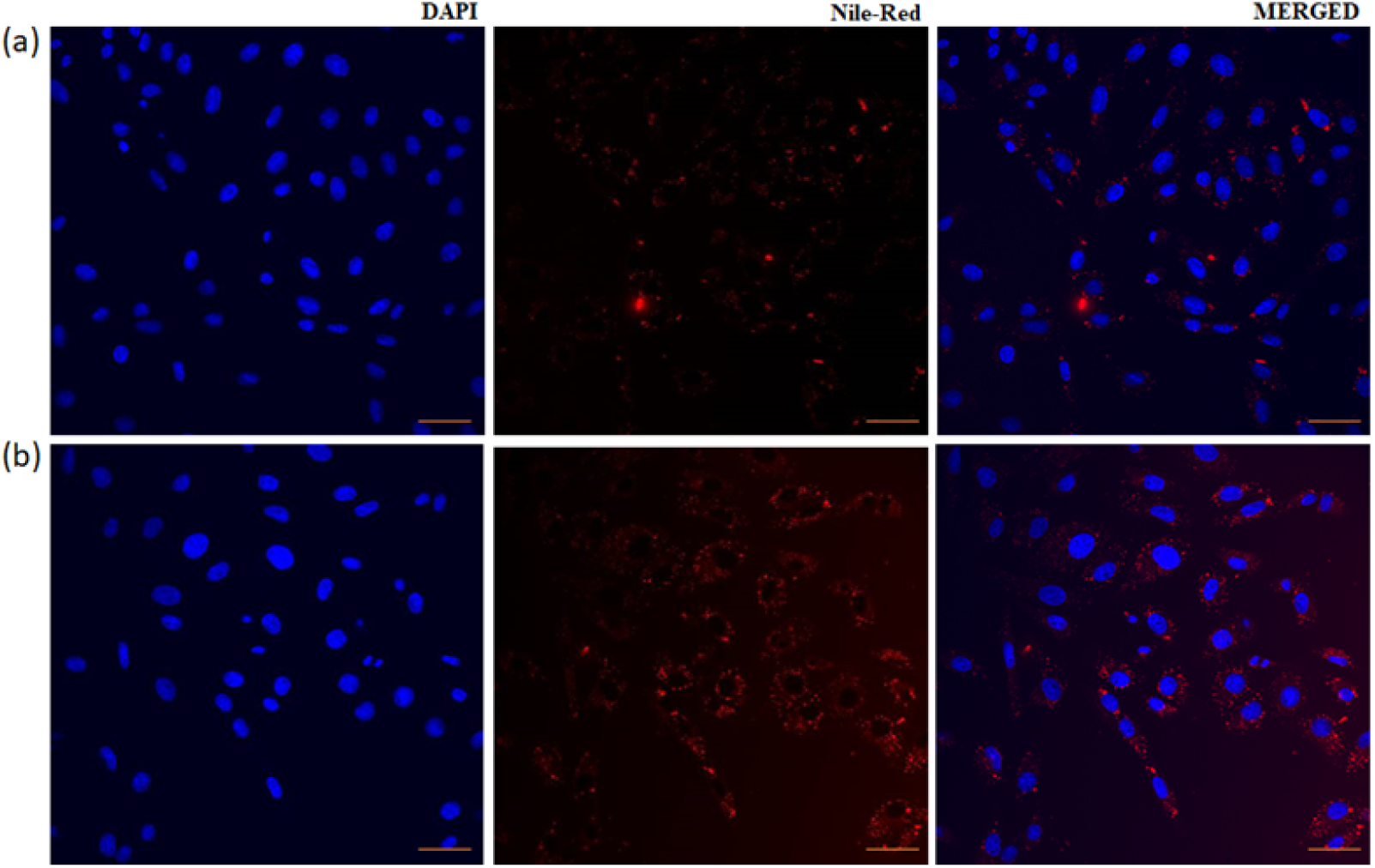
Distribution of lipid droplets in the cell in the presence and absence of 25HC. Panel (a) Less number of lipid droplets proclaim more cholesterol in the cell membrane in the absence of 25HC. Panel (b) the presence of 25HC marks depletion of cholesterol indicating more lipid droplets, stained with Nile-red. Scale bar is 40µm.

Therefore, it is likely that 25 HC might impair the endocytosis by JEV by depleting accessible cholesterol at the plasma membrane, thereby altering the preexisting domain or raft organizations at the plasma membrane. In addition, 25 HC may alter the organization and dynamics (e.g., the fluidity) of the plasma membrane due to the presence of the additional OH group.^10^. However, our time-point data indicate that 25 HC impacts multiple stages of the JEV life cycle, warranting further investigation. Overall, we show that 25 HC is an efficient inhibitor of JEV infection.

25-HC is the product of the interferon-inducible enzyme CH25H. It was earlier reported that CH25H expression is significantly upregulated upon infection by human parainfluenza virus type 3 (HPIV3) and respiratory syncytial virus (RSV), SARS Cov2 and Zika virus.^16^ 25-HC demonstrates a broad range of antiviral activity; however, it’s *in vitro* antiviral activity against JEV has not been reported. Our results show that 25 HC efficiently inhibits JEV infection in Vero cells and warrants further investigation. Antiviral drug development against JEV is rather sluggish and has only limited success. ^17^ Due to the increase in mosquito population, JEV is rapidly spreading to various parts of the world and, therefore there is an urgent unmet need to develop sustainable antiviral strategies against JEV. In this context, our finding may lead to a new treatment against JEV as well as a novel pan-antiviral strategy against flaviviruses.

## Author Contributions

KA carried out experiments, collected and analyzed data, and wrote the first draft; BQ performed the western blot experiments; BB generated and characterized the antibody. SH conceptualized, supervised, analyzed data, wrote, and edited the manuscript secured funding. RKT supervised, edited the manuscript, and secured funding.

## Acknowledgments.

Km. Archana is thankful to UGC for a Senior Research Fellowship and Babita Bohra is thankful to CSIR for a Senior Research Fellowship. We thank the Sophisticated Analytical Instrument Facility (SAIF), CSIR-CDRI, for providing us with the Confocal Microscopy (Intravital facility) and Flow Cytometry facilities. We thank Dr Vikas Agrawal, at the Sanjay Gandhi Postgraduate Institute of Medical Sciences (SGPGIMS), Lucknow for providing the Japanese Encephalitis Virus.

